# Metagenomic data for *Halichondria panicea* from Illumina and Nanopore sequencing and preliminary genome assemblies for the sponge and two microbial symbionts

**DOI:** 10.1101/2021.10.18.464794

**Authors:** Brian W Strehlow, Astrid Schuster, Warren R Francis, Donald E Canfield

**Affiliations:** Department of Biology, University of Southern Denmark, Campusvej 55, Odense M, DK-5230, Denmark

**Author notes:** Denotes corresponding author.

**Keywords:** Metagenome, hologenome, *Halichondria panicea*, Porifera, microbiome

## Abstract

**Objectives:** These data were collected to generate a novel reference metagenome for the sponge *Halichondria panicea* and its microbiome for subsequent differential expression analyses.

**Data description:** These data include raw sequences from four separate sequencing runs of the metagenome of a single individual of *Halichondria panicea* - one Illumina MiSeq (2×300 bp, paired-end) run and three Oxford Nanopore Technologies (ONT) long-read sequencing runs, generating 53.8 and 7.42 Gbp respectively. Comparing assemblies of Illumina, ONT and an Illumina-ONT hybrid revealed the hybrid to be the ‘best’ assembly, comprising 163 Mbp in 63,555 scaffolds (N50: 3,084). This assembly, however, was still highly fragmented and only contained 52% of core metazoan genes (with 77.9% partial genes), so it was also not complete.

However, this sponge is an emerging model species for field and laboratory work, and there is considerable interest in genomic sequencing of this species. Although the resultant assemblies from the data presented here are suboptimal, this data note can inform future studies by providing an estimated genome size and coverage requirements for future sequencing, sharing additional data to potentially improve other suboptimal assemblies of this species, and outlining potential limitations and pitfalls of the combined Illumina and ONT approach to novel genome sequencing.

## Objective

These data were generated to create a reference metagenome for the emerging model sponge species, *Halichondria panicea* and its microbiome. The goal was then to use this reference to study changes in gene expression under different oxygen concentrations in order to understand how this species tolerates hypoxia [see 1]. During the process of data collection, Knobloch et al. [2] generated a reference genome for the dominant microbial symbiont ‘Candidatus Halichondribacter symbioticus’, and the data presented here were not sufficient to construct a suitable reference genome for the sponge, limiting the scope of these data for a full research paper.

Given the considerable interest in *H. panicea* and its widespread distribution, we think that the data provided can inform future experiments, and contribute to a more complete genome later. Finally, by sharing suboptimal data we aimed to identify some potential pitfalls for future genome projects, particularly those of poriferans.

## Data description

### Sample collection and DNA extraction

To limit assembly issues caused by allelic variation, a single individual of *H. panicea* (approximately 1 gram of tissue [wet weight]) was collected from the inlet to Kerteminde Fjord in Denmark (decimal degrees: 55.449808, 10.661299) in 2018 and flash frozen in liquid nitrogen. DNA was extracted using a modified phenol-chloroform extraction (see dx.doi.org/10.17504/protocols.io.yvkfw4w for full protocol). This protocol yielded the highest quality DNA and highest concentrations above 15,000 bp compared to five different extraction protocols (see supplementary material).

In total, nine micrograms of double stranded DNA were extracted and Nanodrop A260/A280 and A260/A230 ratios were 1.79 and 2.17, respectively. The DNA integrity number (DIN) was 1.6, with high concentrations of DNA between 100 and 4,000 base pairs (bp). A smearing pattern in gels was observed for all DNA extractions of *H. panicea* using various protocols (see supplementary material). This pattern could indicate high levels of degradation; however, a substantial amount of DNA was still intact and >15,000 bp long in samples used for sequencing.

### Sequencing

Approximately 1 μg of DNA was sequenced on an Illumina MiSeq sequencer (2×300 bp, paired-end, Illumina, Inc). This run generated 356 million paired-end reads (53.8 Gbp).

The first sequencing run using Oxford Nanopore Technologies (ONT) generated 1.26 million reads (3.4 Gbp, read N50: 2700 bp, longest read: 39,702 bp). For more details on the sequencing methods, see the supplemental material.

Due to a low coverage of Opisthokonta contigs (from the Illumina data) in the nanopore reads, two additional rounds of nanopore sequencing were performed after whole genome amplifications (WGA, see supplementary material), generating 4.021 Gbp from the amplified *H. panicea* DNA. A summary of the public locations of all data generated is shown in Table 1.

**Table 1:** Overview of data files/data sets.

| Label | Name of data file/data set | File types (file extension) | Data repository and identifier (DOI or accession number) |
| --- | --- | --- | --- |
| DNA extraction protocol | Extracting high molecular weight DNA from *Halichondria panicea* (phylum: porifera) | .io | dx.doi.org/10.17504/protocols.io.yvkfw4w |
| Illumina raw sequences lane 1 | 18100FL-02-01-37_S86_L003_R1_001.fastq.gz<br>18100FL-02-01-37_S86_L003_R2_001.fastq.gz | fastq | SRR15711138 (NCBI, SRA) |
| Illumina raw sequences lane 2 | 18100FL-02-01-37_S86_L004_R1_001.fastq.gz<br>18100FL-02-01-37_S86_L004_R2_001.fastq.gz | fastq | SRR15711137 (NCBI, SRA) |
| Nanopore run 1 raw sequences | *H_panicea_ONT_1.fastq* | *fastq* | SRR15711136 (NCBI, SRA) |
| Nanopore run 2 WGA raw sequences | *H_panicea_ONT_2.fastq* | *fastq* | SRR15711135 (NCBI, SRA) |
| Nanopore run 3 - WGA raw sequences | *H_panicea_ONT_3.fastq* | *fastq* | SRR15711134 (NCBI, SRA) |
| Whole metagenome assembly (from illumina sequences) | *JAIOUG01.1.fsa_nt.gz* | *fasta* | JAIOUG000000000, SAMN20669589 (NCBI; GenBank, BioSample accession) |
| *H. panacea* genome assembly (bin 1 from illumina sequences) | *JAIOUD01.1.fsa_nt.gz* | *fasta* | JAIOUD000000000, SAMN20669590 (NCBI; GenBank, BioSample accession) |
| HOC36 bin assembly (from illumina sequences) | *JAIOUE01.1.fsa_nt.gz* | *fasta* | JAIOUE000000000, SAMN20669591 (NCBI; GenBank, BioSample accession) |
| Proteobacteria bin assembly (from illumina sequences) | *JAIOUF01.1.fsa_nt.gz* | *fasta* | JAIOUF000000000, SAMN20669592 (NCBI; GenBank, BioSample accession) |
| Nanopore only metagenome assembly | *JAIOUH01.1.fsa_nt.gz* | *fasta* | JAIOUH000000000, SAMN20669593 (NCBI; GenBank, BioSample accession) |
| Hybrid nanopore-illumina assembly | *JAIOUI01.1.fsa_nt.gz* | *fasta* | JAIOUI000000000, SAMN20669594 (NCBI; GenBank, BioSample accession) |
| Supplementary material | *H_panicea_metagenome_supplemental* | *pdf* | bioRxiv |

### Genome assembly and annotation

#### Illumina metagenome assembly

Full details of quality control, binning, assembly and annotation of the metagenome are in the supplementary material. Three bins were produced including: 1) a large Opisthokonta bin, which was labeled as the sponge bin; 2) a bin for a *Gammaproteobacteria* of the order ‘HOC46’; and 3) a *Proteobacteria* bin (Table 1, 2). The sponge bin was highly fragmented (63,555 scaffolds) and contained only 51.57% of core metazoan genes (with 77.46 % partial matches, Table 2) measured using BUSCOv5 [3]. More bins could potentially be extracted from these data in the future.

**Table 2:**
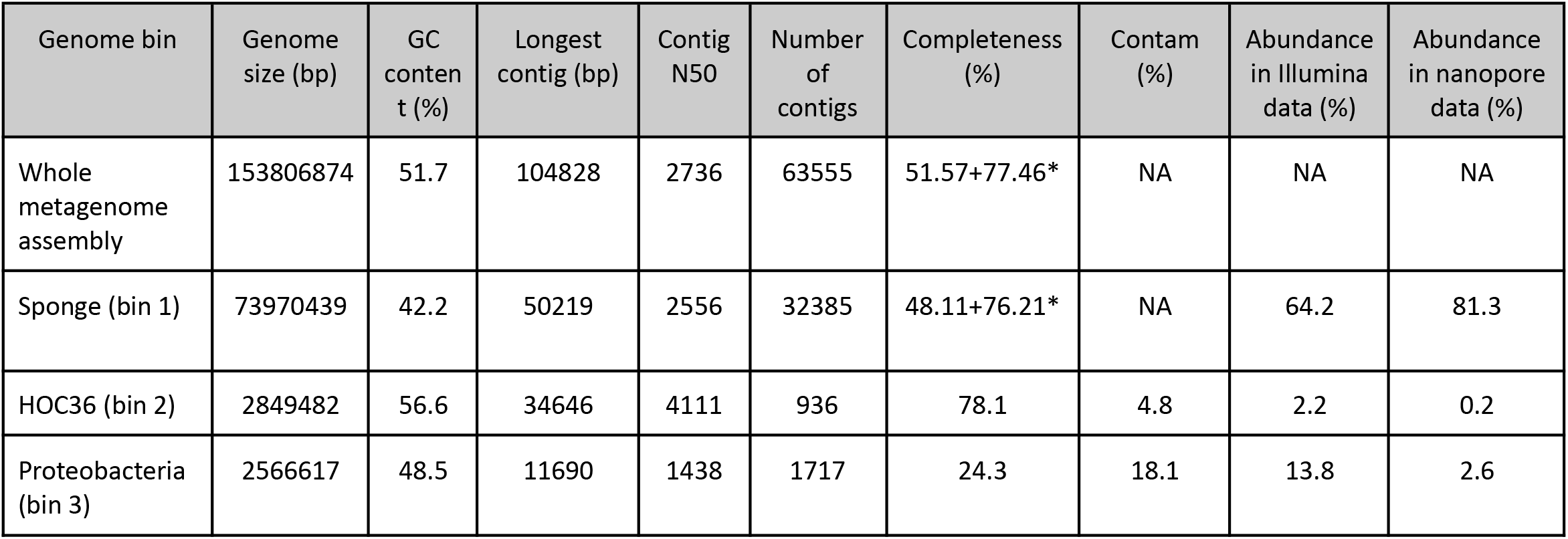
Metagenome statistics from Illumina based assembly. Genome completeness was estimated by CheckM based on the presence of essential single copy genes for prokaryotic bins. For the eukaryotic bin, the percentage of complete and partial core metazoan genes was calculated using BUSCO. *denotes completeness percentage including partial matches. Contamination was estimated by CheckM based on the presence of duplicated single copy genes. Abundance is the relative abundance of each bin compared to the entire metagenome assembly.

| Genome bin | Genome size (bp) | GC content (%) | Longest contig (bp) | Contig N50 | Number of contigs | Completeness (%) | Contam (%) | Abundance in Illumina data (%) | Abundance in nanopore data (%) |
| --- | --- | --- | --- | --- | --- | --- | --- | --- | --- |
| Whole metagenome assembly | 153806874 | 51.7 | 104828 | 2736 | 63555 | 51.57+77.46* | NA | NA | NA |
| Sponge (bin 1) | 73970439 | 42.2 | 50219 | 2556 | 32385 | 48.11+76.21* | NA | 64.2 | 81.3 |
| HOC36 (bin 2) | 2849482 | 56.6 | 34646 | 4111 | 936 | 78.1 | 4.8 | 2.2 | 0.2 |
| Proteobacteria (bin 3) | 2566617 | 48.5 | 11690 | 1438 | 1717 | 24.3 | 18.1 | 13.8 | 2.6 |

The two bacterial genome bins were annotated using PROKKA v. 1.14 [4], and their completeness was estimated with CheckM [5] (see supplemental for more information about these two bins).

#### ONT and hybrid assemblies

Two additional metagenome assemblies were made using 1) ONT data from all three sequencing runs and 2) a combination of Illumina and ONT data. The second ONT sequencing run (following WGA) had high percentages of contamination (8%) and chimerism (5-10%). These ONT data were polished and filtered to remove these errors as described in the supplementary material. A summary of the nanopore-only metagenome assembly is shown in Table 3.

**Table 3.**
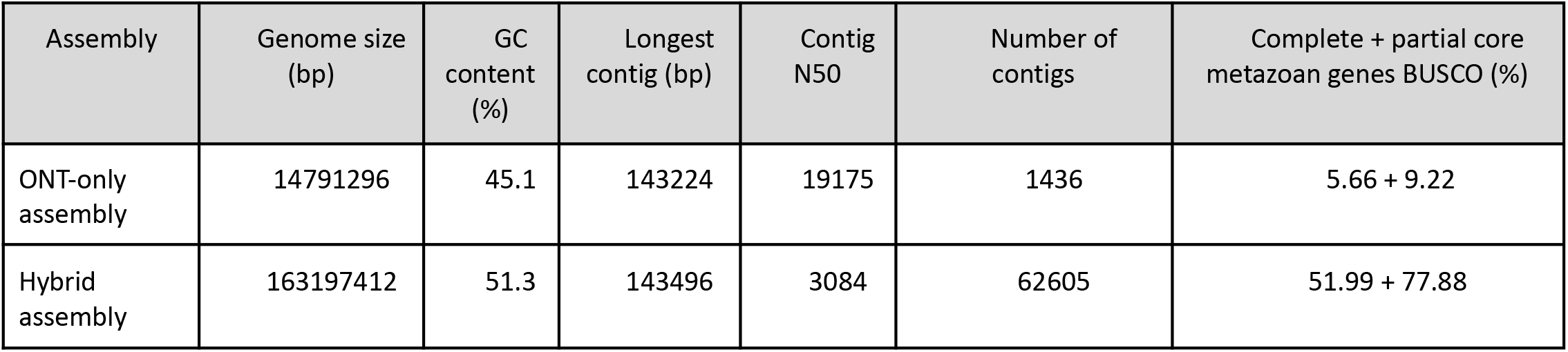
Genome assemblies with nanopore and hybrid (Illumina-nanopore) data.

The ONT and Illumina (Table2) sponge assemblies were merged to create a hybrid metagenome using Flye v.2.6 [6] (Table 3).

### Limitations

Although the incorporation of long read nanopore data in the hybrid assembly did slightly increase the metagenome N50 and decrease the number of scaffolds in the assembly, the genome was still highly fragmented. A major limitation in sponge genomics that is often discussed but rarely written about is the difficulty in extracting high quality, high molecular weight DNA. This difficulty was likely either a result of some innate, highly efficient DNA degradation pathway in *H. panicea* or indicated the presence of DNA and/or degradation pathways from associated microorganisms or secondary metabolites. Obtaining high molecular weight DNA is paramount for successful long-read sequencing as well as genome assembly downstream regardless of sequencing technique. ONT sequencing can selectively sequence smaller DNA fragments if they are present. Additionally, microbial diversity within the metagenome and potential genetic variation caused by diploidy could also have limited genomic assembly.

This note represents the first attempt to sequence a sponge genome using Nanopore and Illumina sequencing, so improved genomic DNA recovery might validate this combination of methods, although it is unclear how DNA recovery could be improved. However, at least 9,000 Mbp long reads need to be generated. Similarly the coverage of ONT reads would need to be increased to ~70x to permit a better assembly. Additionally, WGA should be used with caution due to the high rates of chimerism and contamination throughout the process. Improving coverage would also improve the assembly of prokaryotic genomes in the metagenome.

Recently, the generation of a near-chromosome level scaffolded genome assembly for the sponge *Ephydatia muelleri* was accomplished using PacBio, Chicago, and Dovetail Hi-C libraries sequenced to ~1490x coverage [7]. This sequencing method may therefore be the best for *de novo* genomes. The use of a sponge with limited microbial ‘contamination’ might also be critical for smooth genome assembly, although this effectively limits metagenomic projects. Finally, the use of a single haploid cell, like a sperm or egg cell, could improve future genome assembly performance by limiting allelic variation. However, single cell genomics could be limited by the amount and quality of DNA that can be isolated from a single cell.

## Abbreviations

bp: base pair(s)
ONT: Oxford Nanopore Technologies
WGA: whole genome amplification

## Declarations

### Ethics approval and consent to participate

Not applicable.

### Consent for publication

Not applicable.

### Availability of data and materials

The data described in this Data Note can be freely and openly accessed on NCBI under the project PRJNA753045.

### Authors’ contributions

All authors participated in the conception and planning of the project and reviewed and contributed to drafts of the paper. BWS collected and analyzed the data, contributed reagents/materials/analysis tools, prepared the first draft of the paper, and prepared figures and tables. AS collected the data and contributed reagents/materials/analysis tools. WRF analyzed the data. DEC contributed reagents/materials/analysis tools.

## Acknowledgements

Special thanks to DNASense and Rasmus Dam Wollenberg for the sequencing and downstream support.

