## Supplementary material for "Metagenomic data for *Halichondria panicea* from Illumina and Nanopore sequencing and preliminary genome assemblies for the sponge and two microbial symbionts"

### Sample collection and DNA extraction

DNA quality and quantity were verified using Nanodrop NP-1000 (Thermo Scientific, USA) and double stranded DNA (dsDNA) quantity was verified using a Qubit 4 Fluorometer (Thermo Scientific, USA), using the high sensitivity assay. The DNA length distribution was assessed using a Tapestation 2200 (Agilent Technologies, USA).

DNA was extracted using five other protocols but the linked protocols.io method (below) was judged to be the best. The other extraction protocols included: phenol-chloroform extraction [1], TRIzol (Invitrogen, USA), DNeasy Blood & Tissue Kit (Qiagen, Germany), DNeasy PowerSoil Pro Kit (Qiagen, USA), and Fast DNA Spin Kit for Soil (MP BIO, USA). Test extractions were performed on freeze dried and fresh tissue for comparison using multiple individuals. The phenol-chloroform protocol (dx.doi.org/10.17504/protocols.io.yvkfw4w) yielded the highest concentrations of DNA (>15,000 bp).

### Sequencing

Illumina libraries were prepared using a MiSeq Reagent kit v3 with 600 cycles (Illumina, USA) according to the manufacturer's protocol, and the sample was sequenced across two lanes.

Oxford Nanopore Technologies (ONT) sequencing was performed following the ONT 1D genomic DNA by ligation protocol (SQK-LSK109), using 1 µg of DNA with the L fragment buffer to enrich DNA fragments over 3 kb. CleanNGS magnetic beads (CleanNA, Netherlands) were used, but otherwise the ONT protocol (SQK-LSK109) was followed. The library was sequenced on a MinION sequencer using a FLO-MINSP6 flow cell (ONT, UK). After sequencing, base calling for 1D reads was performed using Guppy (v2.3.5, ONT).

The second two rounds of ONT sequencing were performed following whole genome amplification (WGA) using the REPLI-g Mini Kit (Qiagen, Germany) for 5 µl of DNA. In the first of these, DNA was amplified using WGA and then sequenced as above. In the second round of post-WGA sequencing, for reference and to determine possible chimerism caused by WGA, the *H. panicea* amplified DNA was sequenced along with samples from *E. coli* (unamplified), *E. coli* (WGA), and water (WGA) using the ONT SQK-LSK9 barcoding protocol with the EXPNBD104 kit and the barcoded DNA from these four samples

DNA was pooled. The pool generated 6 Gbp in total for the multiplex. Sequences were basecalled as above and demultiplexed. All ONT runs were performed on FLO-MIN106D flow cells.

### Genome assembly and annotation

#### Illumina metagenome assembly

Adapter sequences were trimmed using cutadapt v. 1.16 [2]. The trimmed reads were assembled into a metagenome using MEGAHIT v. 1.1.3 [3]. This metagenome is summarized in Table 2. To generate coverage files for metagenomic binning, reads were mapped to the assembly using Mini.map2 v. 2.12-r827 [4] . The metagenome was BLASTed v. 2.2.28+ [5] against the SILVA database to identify rRNA genes. These genes were classified using SINA v. 1.2.11 [6] with default settings.

Three metagenome bins were created using the Mmgenome2 package v. 2.0.12 [7] in R v. 3.5.1 based on kmer based tSNE (T-distributed Stochastic Neighbor Embedding).

The closest relative of the HOC36 genome (bin 2) was an uncultured *Proteobacterium* cloned from the sponge *Hymeniacidon heliophila* (94.7% identity from a BLAST of the 16S rRNA gene against NCBI nr database, GenBank: KT880353.1). No rRNA sequence was retrieved for bin 3, but based on all non-rRNA genes, this bin likely belonged to a *Proteobacteria*. Based on their completeness, bins 2 and 3 were expected to have relative abundances of 2.7% and 37.3%, respectively.

#### ONT and hybrid assemblies

These contaminants were filtered out using What's In My Pot (WIMP, Oxford Nanopore) and NanoPlot v.1.27 [8]. The WGA dataset was further filtered for a minimum read length of 2,000 bp and a max base calling error rate of 20%. These filtered reads were combined with filtered reads from the other ONT runs and assembled using Flye v.2.6 [9]. The assembly was then polished using Racon v.1.4.3 [10], Medaka v.0.9.2, with ONT data and again with Racon (with Illumina data). Contigs shorter than 200 bp were trimmed from assemblies to meet NCBI submission requirements.

7. Karst SM, Kirkegaard RH, Albertsen M. mmgenome: a toolbox for reproducible genome extraction from metagenomes. doi:10.1101/059121.

8. De Coster W, D'Hert S, Schultz DT, Cruts M, Van Broeckhoven C. NanoPack: visualizing and processing long-read sequencing data. Bioinformatics. 2018;34:2666–9.

9. Kolmogorov M, Yuan J, Lin Y, Pevzner PA. Assembly of long, error-prone reads using repeat graphs. Nat Biotechnol. 2019;37:540–6.

10. Vaser R, Sović I, Nagarajan N, Šikić M. Fast and accurate de novo genome assembly from long uncorrected reads. Genome Res. 2017;27:737–46.
